# Chromosome-scale genome assembly of *Camellia crapnelliana* provides insights into the fatty acid biosynthesis

**DOI:** 10.1101/2024.01.07.574508

**Authors:** Fen Zhang, Li-ying Feng, Pei-fan Lin, Ju-jin Jia, Li-zhi Gao

**Affiliations:** Engineering Research Center for Selecting and Breeding New Tropical Crop Varieties, Ministry of Education; Tropical Biodiversity and Genomics Research Center, Hainan University, Haikou 570228, China

## Abstract

*Camellia crapnelliana* Tutch., belonging to the Theaceae family, is an excellent landscape tree species with high ornamental value. It is particularly an important woody oil-bearing plant with high ecological, economic, and medicinal values. Here, we first report the chromosome-scale reference genome of *C. crapnelliana* with integrated technologies of SMRT, Hi-C and Illumina sequencing platforms. The genome assembly had a total length of ∼2.94 Gb with contig N50 of ∼67.5 Mb, and ∼96.34% of contigs were assigned to 15 chromosomes. In total, we predicted 37,390 protein-coding genes, ∼99.00% of which were functionally annotated. Comparative genomic analysis showed that the *C. crapnelliana* genome underwent a whole-genome duplication event shared across the *Camellia* species and an γ -WGT event that was shared by all core eudicot plants. Furthermore, we identified the major genes involved in the biosynthesis of oleic acids and terpenoids in *C. crapnelliana*. The chromosome-scale genome of *C. crapnelliana* will become valuable resources for understanding the genetic basis of the fatty acid biosynthesis, and greatly facilitate the exploration and conservation of *C. crapnelliana*.

## Background & Summary

As one of the four largest woody oil plants in the world, oil-tea camellia trees are a collective term for a group of *Camellia* species of highly economic values^1^. In China, oil-tea camellia trees have a long history of cultivation, which are mainly distributed in south of the lower reaches of the Yangtze River^2,3^. There are approximately 50 species of such oil-tea camellia trees, which belong to the family Theaceae^4^. *C. oleifera*, *C. chekiangoleosa*, *C. crapnelliana* and *C. vietnamensis*^1,3^ are commonly cultivated. They are woody, oil-bearing tree species with a high content of seed oil that is widely processed into skin and health care products and especially edible oil^4^. Camellia oil is remarkably rich in polyphenols, saponins, and other healthy components and free of cholesterol, erucic acid, and other harmful components^5^. Thus, the oil has extremely high nutritional and health-beneficial values and has strong market competitiveness and wide market prospects^6^. The content of unsaturated fatty acids in the edible oil is quite high, reaching approximately 90%, and the content of oleic acid can be approximately 87%^5^. Tea oil is therefore referred to as “Oriental olive oil”^7^, which has both health-beneficial and medicinal values^8^.

Among these oil-tea camellia species, *C. crapnelliana*, which belongs to Sect. Furfuracea and is naturally distributed in Hong Kong, southern Guangxi, northern Fujian, southern Zhejiang and Jiangxi provinces, China, was listed as China’s second-class protected plant and recorded in the China Plant Red Data Book (CPRDB) as early as 1992^9^. As an excellent garden greening species with the largest flowers and fruits (**Fig. 1a, b, c**) in the genus *Camellia*, it has great potential for the industrial development as an oilseed plant^10^. Most recently, several chromosome-level tea tree genomes became publicly available^11–16^, oil-tea camellia tree genome information is still quite limited^11–16^. Many efforts have been put on the regulation of the fatty acid biosynthesis in many plants^17–29^, but the molecular basis of the fatty acid biosynthesis remains unclear in *C. crapnelliana*. In-depth understanding about the molecular basis and evolution of the fatty acid biosynthesis in *C. crapnelliana* largely rely on a high-quality reference genome.

**Fig. 1.**
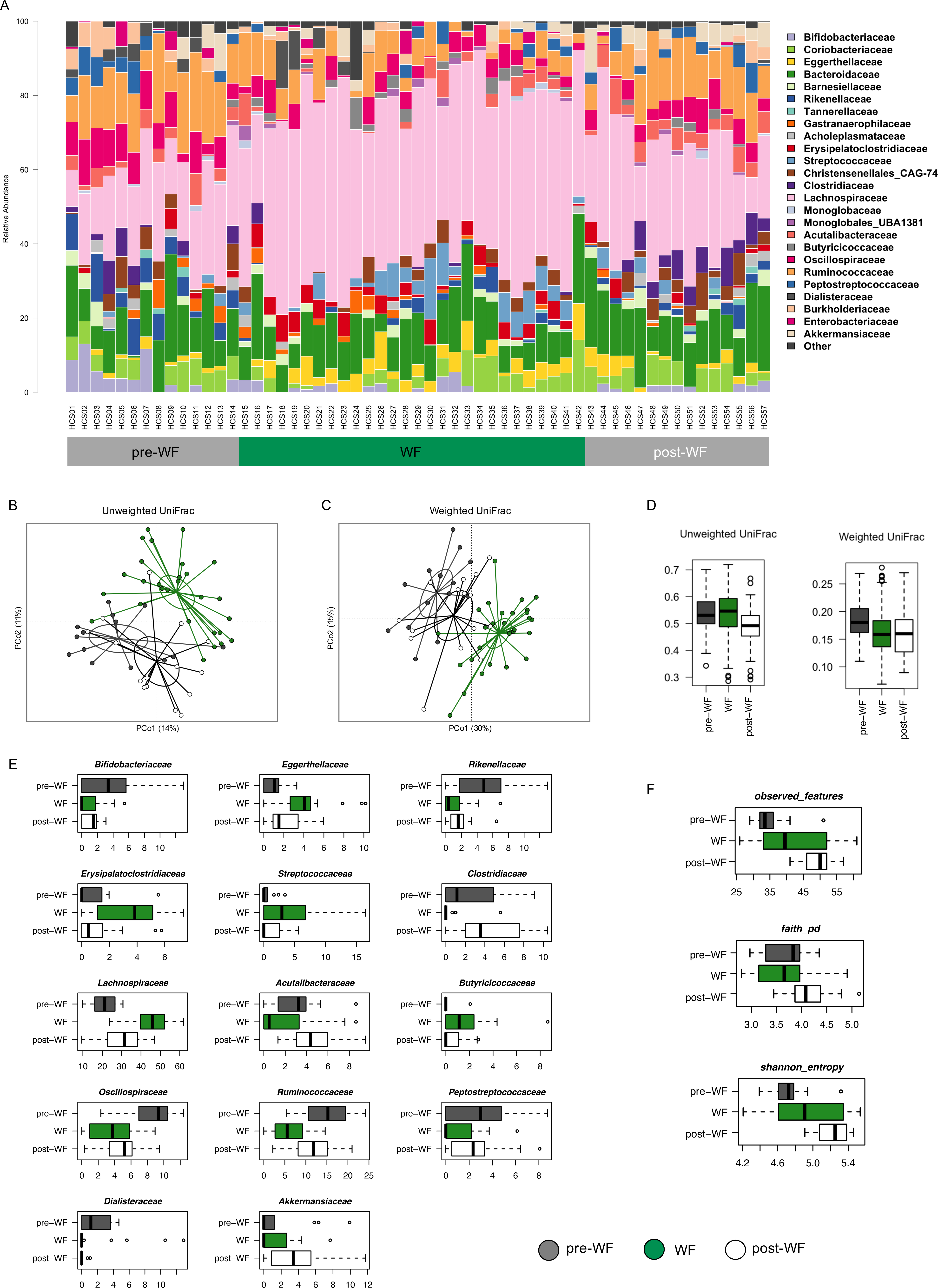
Summary of genome assembly and plant features of *C. crapnelliana*. Tree plant (**a**), flower (**b**), fruit (**c**) and genome assembly statistics (**d**) of *C. crapnelliana*.

In this study, we constructed and annotated a high-quality chromosome-level reference genome of *C. crapnelliana* using integrated sequencing data (∼71×PacBio HiFi reads and ∼140×Hi-C reads) (**Fig. 2**). *K-mer* analysis showed that the genome size of *C. crapnelliana* was estimated to be ∼3.055Gb, with a repeat sequence proportion of 76.76% (**Supplementary Table S2**). The final assembled genome was ∼2.94Gb, with contig N50 of ∼67.50Mb (**Fig. 1d**). Based on the karyotype of the species (2n=30)^30^, approximately ∼96.34% of the contig reads were anchored to 15 pseudochromosomes. A total of 37,390 protein-coding genes were predicted, of which 99.00% were functionally annotated. In addition, 176 miRNAs, 7,988 rRNAs, 857 tRNAs, and 485 snRNAs in the *C. crapnelliana* genome were annotated. Comparative genomic analysis suggested rapid evolution of gene families in the *C. crapnelliana* genome, evidenced by a remarkable expansion of gene families as well as the innovation of a number of unique gene families. Results also revealed that the *C. crapnelliana* genome underwent a whole-genome duplication event shared by all *Camellia* species. we particularly identified the candidate genes associated with the fatty acid biosynthesis in *C. crapnelliana*. The high-quality chromosome-level genome assembly of this oil-tea camellia species will greatly help to enhance the functional analysis and novel genes towards oil quality and yield improvement, and augment wild resources conservation and utilization in the future.

**Fig. 2.**
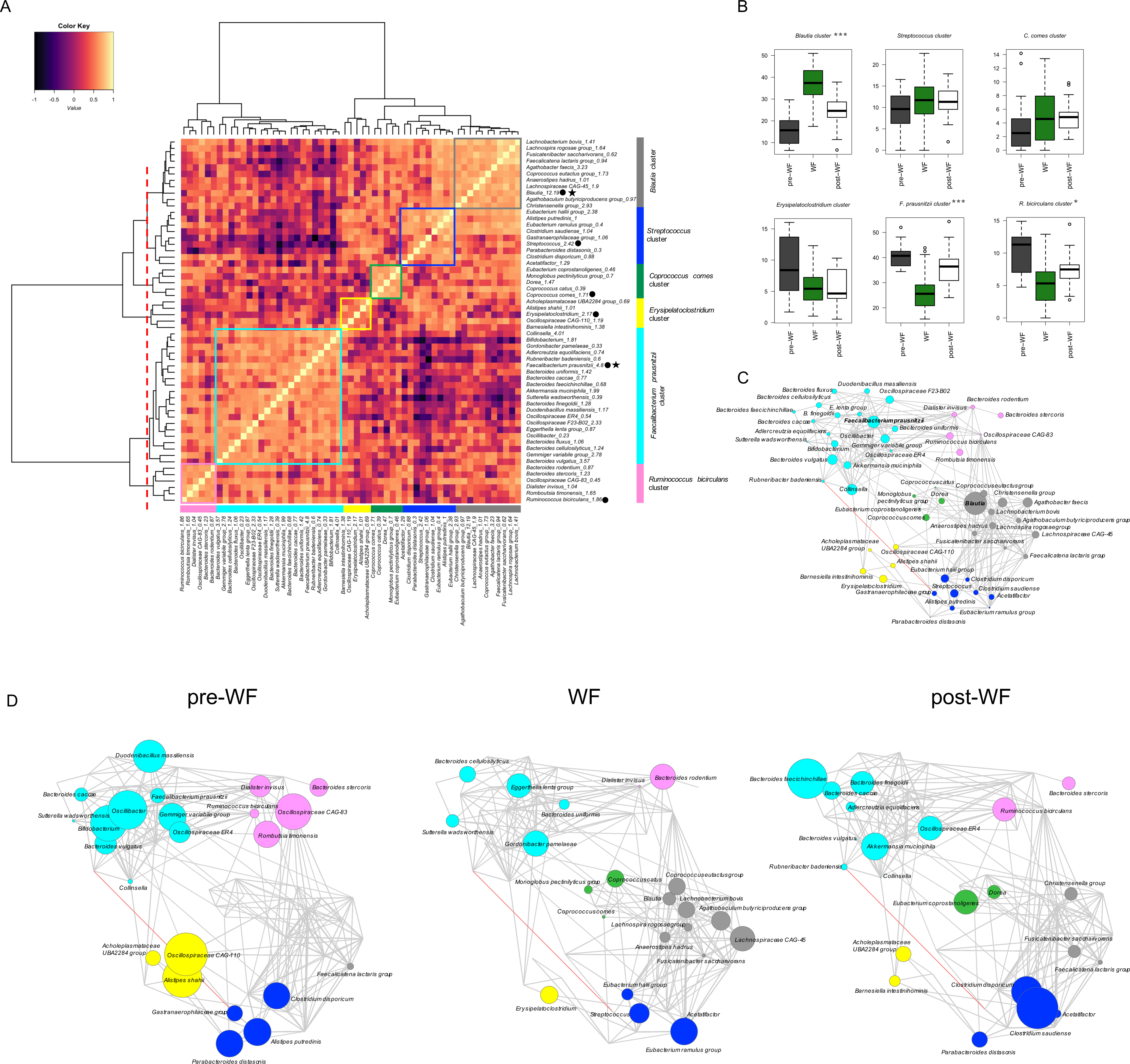
Overview of features of the *C. crapnelliana* genome. The outermost layer represents 15 pseudo-chromosomes of the *C. crapnelliana* genome (scale mark =1Mb), and the second to seventh circles represent the density of protein-coding genes, repeat sequence density, GC content, total TEs, *Gypsy*-like element distribution, and *Copia*-like element distribution. The innermost track indicates genomic synteny among the chromosomes.

## Methods

### Plant materials, sample collection, and sequencing

For genomic DNA extraction, young healthy leaves of *C. crapnelliana* were collected from South China National Botanical Garden in Guangdong Province, China (**Fig. 1b**). Sampled leaves were immediately flash-frozen in liquid nitrogen and stored at −80 °C until further use. High molecular weight genomic DNAs (gDNAs) were extracted from leaves using improved CTAB method^31^ and evaluated using NanoDrop One spectrophotometer (NanoDrop Technologies, Wilmington, DE) and Qubit 3.0 Fluorometer (Life Technologies, Carlsbad, CA, USA). For the genome survey, the paired-end (PE 150bp) library was generated using the Illumina TruSeq DNA Nano Preparation Kit (Illumina, San Diego, CA, USA), and the library was sequenced on an Illumina HiSeq 2500 platform following the manufacturer’s instructions. As a result of Illumina sequencing, we obtained ∼173.51Gb of Illumina paired-end reads (**Supplementary Table S1**). The Pacbio HiFi sequencing was then performed on the PacBio Sequel LJ platform (Pacific Biosciences, CA, USA), according to the manufacturer’s instructions. We obtained ∼212.87Gb HiFi reads with an average read length of ∼19,232.96 bp, which covered about 71× of the *C*. *crapnelliana* genome (**Supplementary Table S1**). For Hi-C sequencing, formaldehyde was used for crosslinking the fresh leaves, and the crosslinking reaction was terminated using glycine solution. Subsequently, the Hi-C library was constructed based on the instructions and sequenced on the Illumina platform (Annoroad Gene Technology Co., Ltd), and ∼429.88Gb raw reads were generated (**Supplementary Table S1**). The young leaves, flowers, young shoots, and seed kernels were collected for transcriptome sequencing. These tissue samples were rinsed using ddH_2_O and stored at −80°C until use after snap-freeze using liquid nitrogen with three biological replicates. Total RNA extraction was performed using the RNeasy Plant Mini Kit (Qiagen, Hilden, Germany). A cDNA library was built following the instructions, followed by paired-end sequencing on the NovaSeq platform (Illumina). A total of ∼30.00 Gb RNA-seq reads were obtained to assist the subsequent analysis of the *C. crapnelliana* genome.

### Chromosome-level genome assembly

Genome size of *C*. *crapnelliana* was estimated from Hi-C data using *k-mer* frequency analysis. Jellyfish v2.3.0^32^ was first applied to extracting and counting canonical *k*-mer at k=21. Subsequently, findGSE v1.94^33^ was used to estimate the genome size from *k-mer* count data with parameters of “-k=21”. As a result, we estimated the genome size of *C. crapnelliana* to be ∼3.055Gb (**Supplementary Table S2**). The PacBio HiFi reads were *de novo* assembled by using hifiasm v0.16.1^34^ with default parameters. The genome assembly had a total size of ∼2.94Gb, containing 816 contigs with N50 sizes of 67.5Mb (**Supplementary Table S3**). The cleaned Hi-C reads were mapped to the corresponding contigs using Juicer v1.9.9^35^. The unique mapped reads were taken as input for 3D-DNA pipeline v180114^36^ with parameters “-r 0” and then sorted and corrected manually using JuicerBox v1.11.08^37^. The fifteen pseudochromosomes were identified by distinct interaction signals in the Hi-C interaction heatmap (**Supplementary Fig. S1**), and the final assembled genome length was ∼2.94Gb (**Fig. 1d, 2**), with a scaffold N50 of ∼67.50Mb, containing ∼96.34% of the assembled contigs for *C. crapnelliana* (**Supplementary Table S4**), accounting for ∼96.34% of the estimated genome size based on the *k-mer* analysis (**Supplementary Table S2**).

### Genome annotation and functional prediction

The repetitive elements in the *C. crapnelliana* genome were identified by combining *de novo* and homology-based approaches. Tandem repeat sequences were annotated using Tandem Repeat Finder (TRF v4.09)^38^ with default parameters. A total of six types (mono-to hexa-nucleotides) of simple sequence repeats (SSRs) were identified using the MISA (MIcroSAtellite)^39^ identification tool with default parameters. For *de novo*-based searches, RepeatModeler v2.0.2a^40^, LTR_FINDER v1.07^41^, LTRharvest v1.5.9^42^, and LTR_retriever v2.9.1^43^ were applied for constructing *de novo* repeat libraries, by which RepeatMasker v4.1.3-p1^44^ was employed to detect repeat sequences. For homology-based searches, we employed RepeatMasker v4.1.3-p1^44^ against a known repeat library, Repbase v.19.06^45^. As a result, a total of ∼2.44Gb of repetitive elements occupying ∼82.87% of the *C. crapnelliana* genome were annotated (**Fig. 2**; **Supplementary Table S7**). Most of these repeats were long terminal repeat (LTR) retrotransposons (∼63.24%) of the genome; **Supplementary Table S7**). The DNA, LINE, and SINE classes accounted for ∼10.84%, ∼4.19%, and ∼0.13% of the genome, respectively (**Fig. 2**; **Supplementary Table S7**). Additionally, tRNAscan-SE v2.0^46^ software was used to predict tRNA genes. The rRNA, miRNA, and snRNA were predicted using INFERNAL (v1.1.2)^47^ software through searches against the Rfam database v9.1^48^. Finally, we annotated 176 miRNAs, 7,988 rRNAs, 857 tRNAs, and 485 snRNAs in the *C*. *crapnelliana* genome (**Supplementary Table S8**).

To annotate protein-coding genes in the *C*. *crapnelliana* genome, gene models were obtained by combining the three approaches of *ab initio* gene predictions, homology-based predictions, and transcriptome-based predictions. The *ab initio* prediction was performed by AUGUSTUS v3.3.2^49^, SNAP^50^ v2013-11-29, GeneMark-ES/ET^51^, GlimmerHMM^52^ v3.02. For homology-based prediction, the Exonerate^53^ v2.2.0 program was used to search against the protein sequences of *Actinidia chinensis*^54^, *Arabidopsis thaliana*^55^, *Beta vulgaris*^56^, *C. oleifera*^26^, DASZ^14^, *C. sinensis* var. *assamica* YK^11^, *Olea europaea*^57^, *C. chekiangoleosa*^58^, *C. lanceoleosa*^59^, *Vitis vinifera*^60,61^, and *Oryza sativa*^55^ genomes. For transcriptome-based prediction, Trinity v2.15.1^62^ was used for assembling transcripts based on RNA-seq data, and PASA^63^ v2.5.2 software was employed for gene structure prediction based on transcriptome assemblies. Additionally, HISAT2 v2.2.1^64^ was employed for RNA-seq reads mapping onto the genome, and StringTie^65^ v2.2.1 was used for the generation of transcript structure. The assembled transcripts were subsequently used for ORF (open reading frame) prediction using TransDecoder v5.5.0. All predicted gene structures were integrated into a consensus set with EVidenceModeler (EVM v2.0.0)^66^. Finally, 37,390 gene models were predicted after integrating the results of the three aforementioned methods (**Fig. 1a, 2**).

For the functional annotation of protein-coding genes, we aligned the predicted protein-coding gene sequences against public functional databases using BLAST v2.11.0^67^ (e-value < 1e-5), including Swiss-Prot^68^, NR^69^, KEGG, and KOG^70^. Gene Ontology (GO) was performed using InterProScan v5.55-88.0^71,72^(**Supplementary Fig. S2**). As a result, a total of 37,015 protein-coding genes were annotated for *C*. *crapnelliana*, accounting for ∼99.00% of all predicted genes (**Supplementary Table S9**). Predicted gene models were comparable to the fifteen other species in aspects such as gene number, average gene length, average CDS length, average exons per gene, average introns per gene, average exon length, and average intron length (**Supplementary Table S10**).

### Comparative genomic analysis

The protein-coding genes of nine representative species were selected for gene family analysis using OrthoFinder v2.5.4^73^ with default parameters, including *Actinidia chinensis*^54^(ACH), *Arabidopsis thaliana*^55^ (ATH)*, C. chekiangoleosa*^58^ (CCH)*, C. sinensis* var*. assamica*^11^(CSA-YK10)*, C. sinensis* var. *sinensis*^12^ (CSS-BY), *C. oleifera*^26^ (CON)*, Olea europaea*^57,74^ (OEU), *Vitis vinifera*^60,61^ (VVI), and *C. crapnelliana* (CCRA). A total of 107 single-copy orthologues were obtained, and they were aligned using MAFFT v7.505^75^, and then ProTest v3.4.2^76^ was used to find the best model of amino acid replacement in the single-copy alignments. Before the phylogeny construction, Trimal v1.4.rev15^77^ was employed to judiciously eliminate ambiguously aligned positions. The final alignments of orthologous groups were concatenated to build a phylogenetic tree using RAxML v8.2.10^78^ with the ML algorithm and 1,000 bootstrap replicates. Divergence times among the species were estimated using r8s v1.80^79^ based on the TimeTree^80^ website (http://timetree.org/). The phylogenetic tree was visualized by iTOL^81^ online tools (https://itol.embl.de/#). The results showed that *C. crapnelliana* was sister to *C. chekiangoleosa* and *C. oleifera* (**Fig. 3a**), supported by previous studies based on chloroplast and nuclear sequences^82,83^. According to the time-calibrated phylogeny, *C. crapnelliana* diverged from *A. chinensis* at 82.8 million years ago^54,84^.

**Fig. 3.**
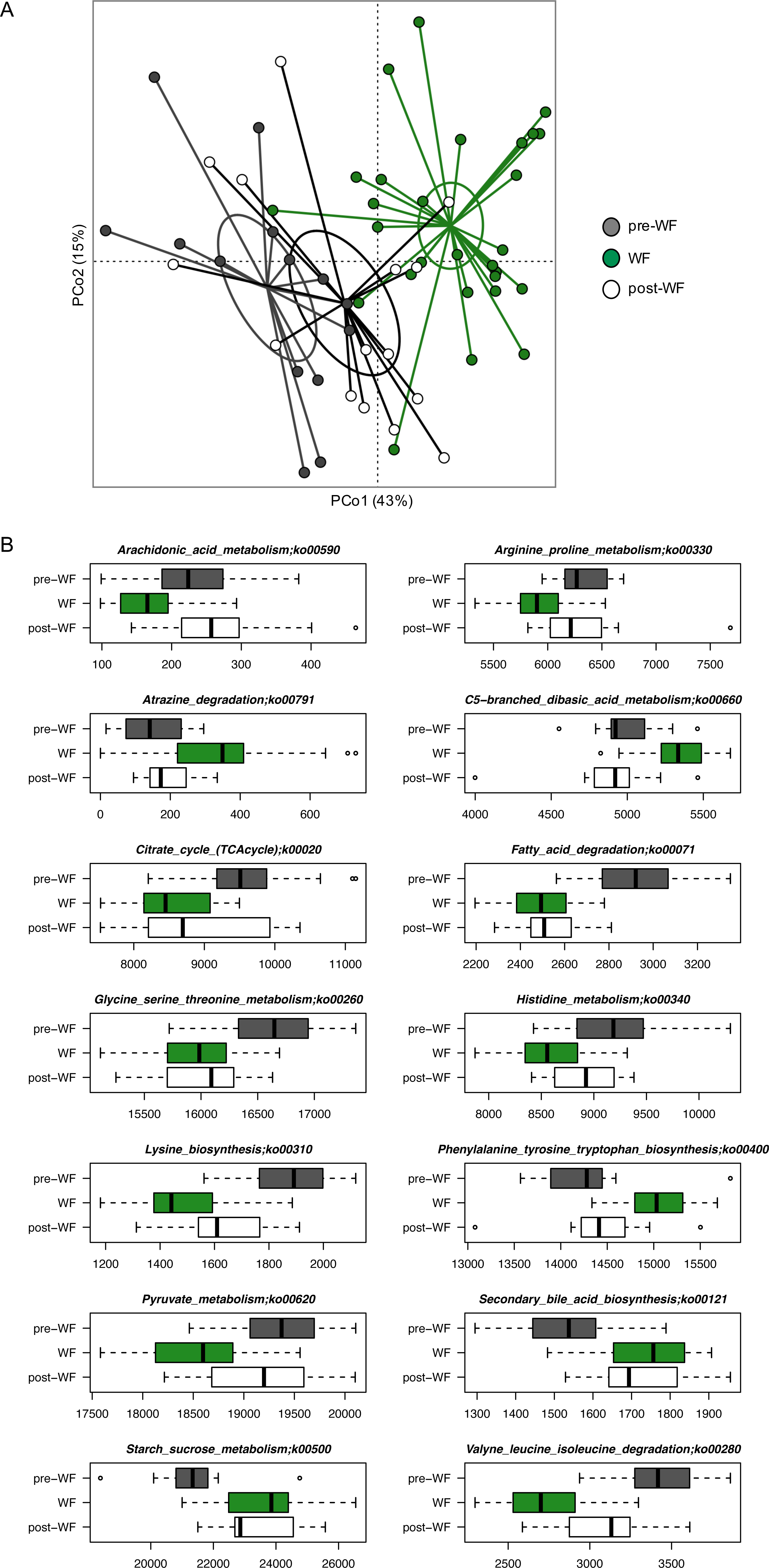
Comparative genomic analysis between *C. crapnelliana* and representative plant species. (**a**) Phylogenetic tree showing the relationships among nine plant species, along with divergence times. Numbers of gene families that underwent contraction (orange) and expansion (green) during evolution are indicated. Different categories of orthologous genes across all species are displayed as stacked bar charts on the right. (**b**) The upset plot shows the results of common and specific gene families in the nine plant genomes. (**c**) KEGG enrichment analysis of the expanded gene families in *C. crapnelliana*.

To shed light on the evolution of gene families, we embraced CAFE^85^ v5.0.0 with default parameters, conceptualizing gene family evolution as a stochastic birth and death process. We further delved into the intricate nuances of gene family evolution. This involved a meticulous analysis of changes in gene family sizes across the phylogenetic tree. These analyses were orchestrated using the R package clusterProfiler. The analysis of gene family evolution showed that *C. crapnelliana* underwent more gene family contraction than expansion (**Fig. 3a**; **Supplementary Table S11**). Among them, 1,127 gene families (*P*-value≤0.01) were significantly expanded in the *C. crapnelliana* genome. Comparative genomic analyses of the nine representative plant genomes also revealed that *C. crapnelliana* possessed 293 species-specific gene families (**Fig. 3b**). Functional enrichment analysis indicated that the expanded gene families were significantly enriched in Gene Ontology (GO) terms of environmental adaptation, biological regulation and defense response (**Fig. 3c**).

### Genome synteny analysis and detection of whole-genome duplication (WGD)

To detect whole genome duplication events in the *C. crapnelliana* and other plant genomes, we used the Whole-Genome Duplication Integrated analysis tool (WGDI v0.6.5)^86^ for the detection of WGDs and intragenomic collinearity as well as *Ks* estimation and peak fitting. The WGD analyses were performed using all paralogous gene pairs. MAFFT v7.520^87^ was employed to conduct sequence alignment. The protein sequence alignment was converted into a codon alignment using PAL2NAL v14. Finally, the *Ka* and *Ks* values were obtained using yn00 v4.10.0 of PAML^88^ with the Nei-Gojobori (NG) method. WGDI was adopted to mark the *Ks* on the syntenic block with different colors. The PeaksFit (−pf), Kspeaks (−kp), and KsFigures (−kf) tools of WGDI were used to illustrate the *Ks* density. JCVI^89^ v1.3.6 was used to draw a collinearity diagram of the current patterns of these species. The *C. crapnelliana* genome exhibited two peaks in the *Ks* density plot (**Fig. 4a, b**). Our results showed that the occurrence of two polyploidization events in the *C. crapnelliana* genome, including the ancient WGT (γ) event that occurred in grape and eudicots^60,61^, the other WGD (β) event shared with *A. chinensis* and other Theaceae species^11,54,84^ (**Fig. 4a, b**). We finally verified the occurrence of two WGD events in the *C. crapnelliana* genome by combining genomic synteny analysis (**Fig. 4d**) and dot plots of *C. crapnelliana* (**Fig. 4c**).

**Fig. 4.**
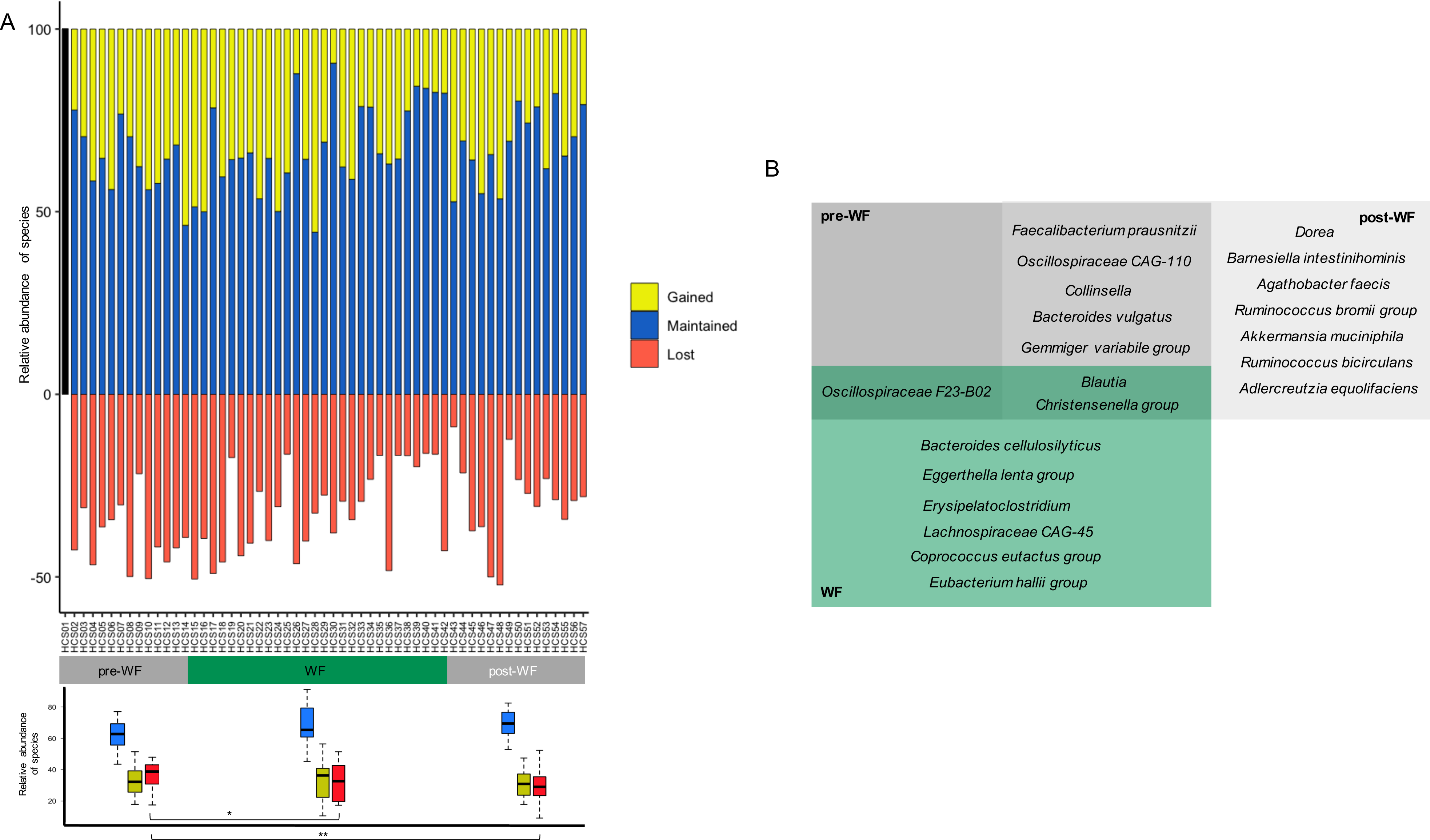
Detection of whole genome duplication (WGD) and genomic synteny analysis between the *C. crapnelliana* and *C. sinensis* genomes. (**a**, **b**) Distribution of *Ks* values in *Actinidia chinensis* (ACH)*, C. sinensis* var*. assamica* (CSA-YK10)*, Vitis vinifera* (VVI), and *C. crapnelliana* (CCRA), which represents the Gaussian fit of the raw *Ks* counts from paralogs. (**c**) Synteny blocks of the *C. crapnelliana* genome. The axes refer to different chromosomes, and genomic synteny blocks represent the WGD event. (**d**) Diagram showing genomic collinearity among the *C. sinensis* var*. sinensis* (CSS-BY), *C. sinensis* var*. assamica* (CSA-YK10), and *C. crapnelliana* (CCRA) genomes.

### Identification of genes involved in the fatty acid biosynthesis

To elucidate the genetic components governing the fatty acid biosynthesis (**Fig. 5a**), we initiated a meticulous retrieval and comparative analysis of relevant genes. The foundation of this analysis lay in the comprehensive aralip database (http://aralip.plantbiology.msu.edu/downloads), housing genes associated with the fatty acid biosynthesis in *A. thaliana*^90^. In a bid to uncover orthologous genes in *C. crapnelliana* and *C. sinensis* var*. assamica*, we initiated a sequence similarity search. Deploying the Blastp v2.11.0+^91^ tool with stringent criteria (e-value ≤ 1e−5, alignment identity ≥ 50%, and Score ≥ 200), we established connections between genes. At the same time, all family genes were predicted by hmmsearch^92^ v3.3.2 with the Pfam accession in **Supplementary Table S12** (e-value < 1e-20). MAFFT v7.505^87^ was employed to perform protein sequence alignments. MEGA11^93^ was used to construct the phylogenetic tree with the ML model (bootstrap set as 1000). We totally identified 57 and 54 the fatty acid biosynthesis-related genes from the *C. crapnelliana* and *C. sinensis* var. *assamica* genomes, respectively (**Fig. 5b**; **Supplementary Table S13**). Furthermore, we identified 114 and 137 triacylglycerol biosynthesis (oil-body) related genes from the *C. crapnelliana* and *C. sinensis* var. *assamica* genomes, respectively (**Supplementary Table S13**).

**Fig. 5.**
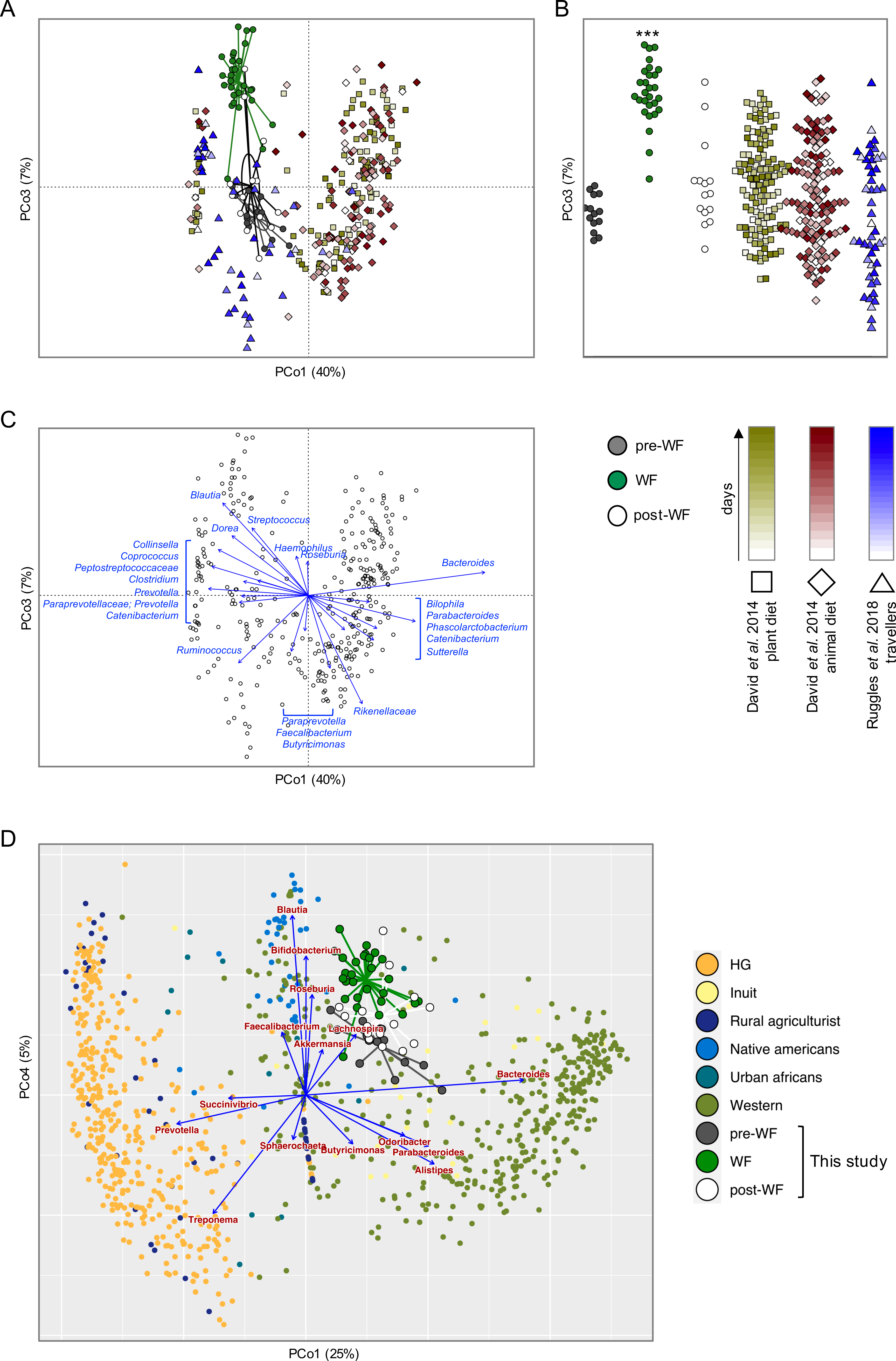
Phylogenetic analysis of candidate genes involved in the fatty acid biosynthesis between *C. crapnelliana*, tea tree and *A. thaliana* genomes. (**a**) The fatty acid biosynthetic pathway. (**b**) Phylogenetic analysis of candidate genes involved in the fatty acid biosynthesis in the *C. sinensis* var*. assamica*, *A. thaliana*, and *C. crapnelliana* genomes.

## Data Records

The MGI short reads, PacBio HiFi long-reads, Hi-C reads, genome assembly and annotation data were deposited in the National Genomics Data Center (NGDC)^94^, Beijing Institute of Genomics, the Chinese Academy of Sciences/China National Center for Bioinformation with BioProject Accession Numbers PRJCA022516^95^. The genome sequencing data were deposited in the Genome Sequence Archive (GSA) of NGDC under Accession Numbers CRA014272^96^. The genome assembly and annotation data were deposited in Genome Assembly Sequences and Annotations (GWH)^97^ of NGDC under accession number GWHERAW00000000^98^. The genome assembly and annotation as well as information of the identified genes involved in the fatty acid biosynthesis were deposited at the figshare database^99^.

## Technical Validation

### Assessment of the genome assembly

The completeness of the assembled genome was evaluated using BWA (v0.7.17)^100^ and Benchmarking Universal Single-Copy Orthologs (BUSCO, v5.4.4)^101^ with the embryophyta_odb10 lineage dataset. Approximately, ∼99.67% of the Illumina short reads were aligned to the genome, of which ∼93.97% of reads were properly mapped. The BUSCO analysis showed that the assembled genome sequences contained 1,600 (∼99.2%) complete BUSCOs, including 1,405 (∼87.1%) single-copy BUSCOs, 195 (∼12.1%) duplicated BUSCOs, and 8 (∼0.5%) fragmented BUSCOs (**Supplementary Table S5**).

### Assessment of the gene annotation

The annotated and integrated proteins were also evaluated using BUSCO v5.4.4^101^ with the lineage dataset embryophyte_odb10. Briefly, the proportion of complete core gene coverage was ∼96.2% (including ∼87.3% single-copy genes and ∼8.9% duplicated genes), and there were only a few fragmented (∼1.4%) and missing (∼2.4%) genes (**Supplementary Table S6**), indicating high-quality annotation of the predicted gene models.

## Code availability

All software and pipelines were executed according to the manual and protocols of the published bioinformatic tools. The version and code/parameters of the software were described in the Methods section.

## Acknowledgements

We would highly appreciate Zi-ting Yu for her assistance to collect samples.

## Author contributions

Li-zhi Gao conceived and designed the study and revised the manuscript. Fen Zhang executed data analysis and drafted the manuscript. Li-ying Feng and Pei-fan Lin collected samples and performed experiments. Ju-jin Jia contributed to data analyses. All authors read, edited, and approved the final manuscript.

## Additional information

**Supplementary Fig. S1.**
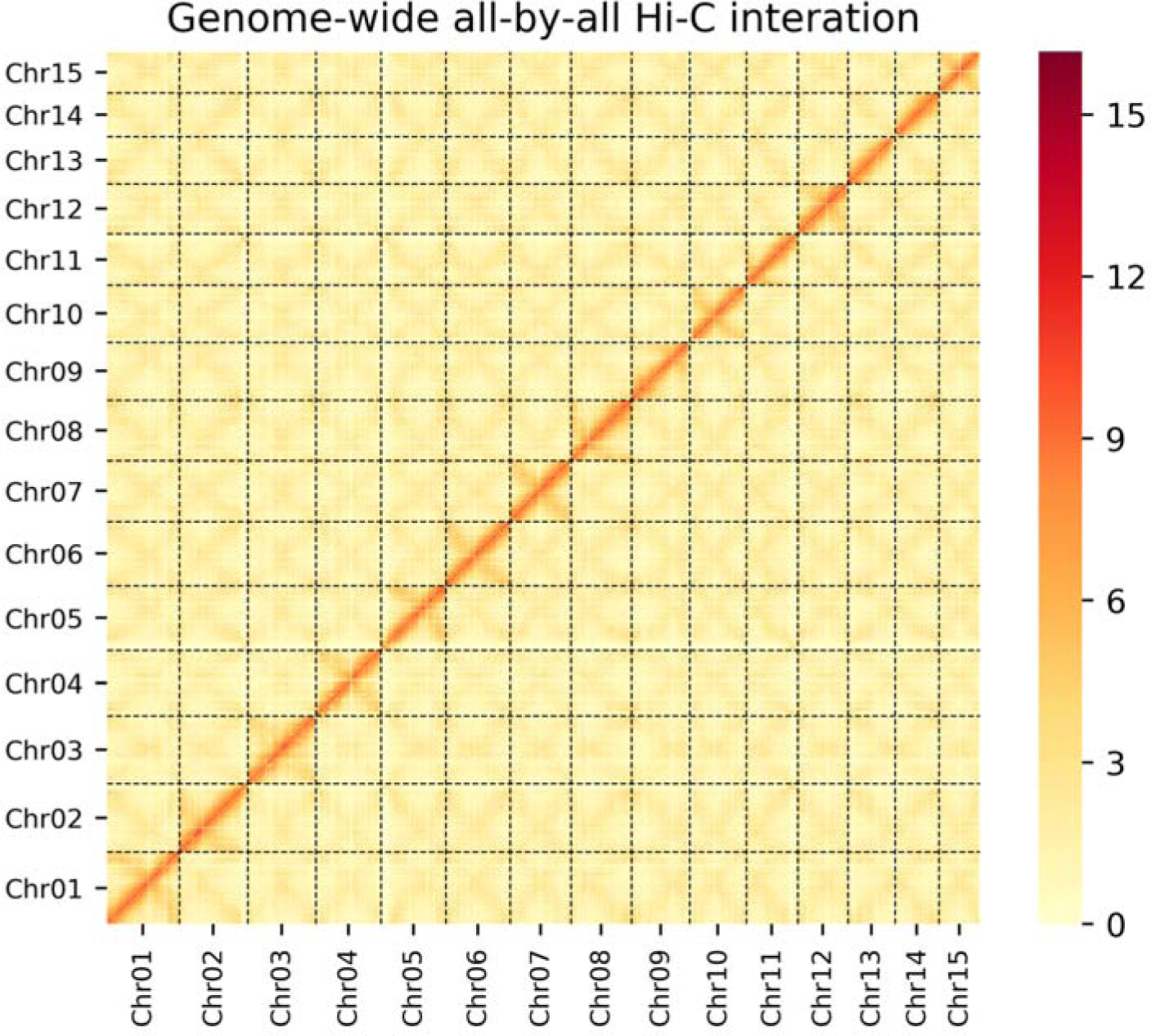
Genome-wide Hi-C interaction map of *C. crapnelliana*. The heat map shows the intensity signals of Hi-C chromosome interaction.

**Supplementary Fig. S2.**
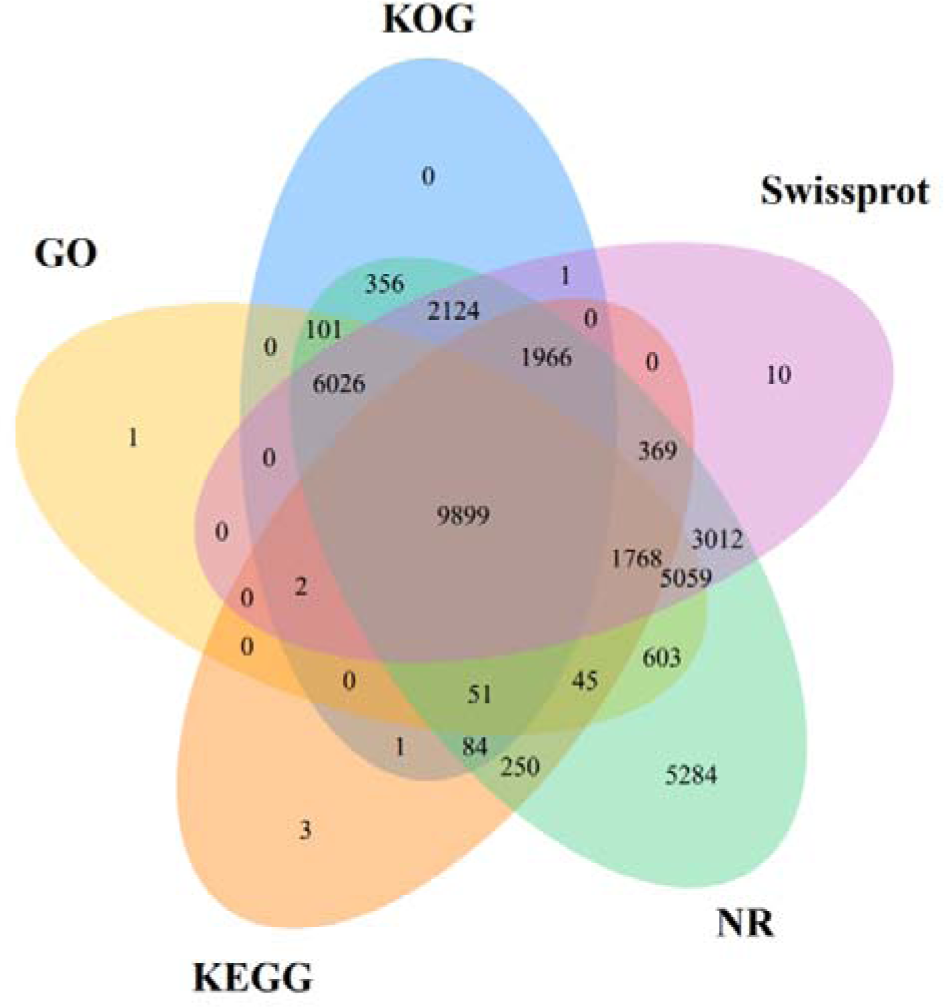
Venn diagram of the number of genes from *C*. *crapnelliana* with homology or functional classification.

