## Supplementary Tables for "Chromosome-scale genome assembly of *Camellia crapnelliana* provides insights into the fatty acid biosynthesis"

**Supplementary Table S1** **Summary of DNA sequencing data for the *C. crapnelliana* genome.**

| **Sequence type** | **Average read length (bp)** | **Raw data (Gb)** | **Raw reads number** | **Depth** | **N50 （bp）** | **Purpose** |
| --- | --- | --- | --- | --- | --- | --- |
| HiFi | 18,993 | 37.05 | 1,950,491 | 12.35× | 19,501 | *de novo* assembly |
| HiFi | 18,071 | 26.35 | 1,458,247 | 8.78× | 18,428 | *de novo* assembly |
| HiFi | 18,857 | 24.96 | 1,323,456 | 8.32× | 19,398 | *de novo* assembly |
| HiFi | 19,152 | 23.75 | 1,240,177 | 7.92× | 19.688 | *de novo* assembly |
| HiFi | 19,774 | 100.76 | 5,095,607 | 33.59× | 20,953 | *de novo* assembly |
| HiC | 150 | 429.88 | 2,865,866,667 | 143.29× | 150 | chromosome-level assembly |
| MGI-Seq | 150 | 173.51 | 1,156,733,333 | 57.84× | 150 | Survey |

**Supplementary Table S2 Survey statistics of the *C. crapnelliana* genome.**

| **Species** | **Total base（Gb）** | ***K-mer*** | ***K-mer* number** | ***K-mer* depth** | **Genome size (Gb)** | **Repeat ratio（%）** |
| --- | --- | --- | --- | --- | --- | --- |
| ***C. crapnelliana*** | 429.88 | 17 | 475,641,075,796 | 157.7 | 3.055 | 76.76 |

**Supplementary Table S3 Statistics of the *C. crapnelliana* genome assembly.**

| **Stat Type** | **Contig Length** | **Contig Number** | **Scaffold Length** | **Scaffold Number** | **Gap Length** | **Gap Number** |
| --- | --- | --- | --- | --- | --- | --- |
| N50 | 67,500,000 | 15 | 197,708,864 | 7 | 100 | 32 |
| N60 | 58,223,584 | 19 | 189,639,866 | 9 | 100 | 38 |
| N70 | 48,000,000 | 25 | 183,972,917 | 10 | 100 | 45 |
| N80 | 36,715,753 | 32 | 161,953,748 | 12 | 100 | 51 |
| N90 | 16,141,345 | 45 | 142,359,996 | 14 | 100 | 57 |
| Longest | 175,800,000 | 1 | 234,637,713 | 1 | 100 | 63 |
| Total | 2,944,514,837 | 816 | 2,944,521,137 | 753 | 6,300 | 63 |
| Length>=1kb | 2,944,514,837 | 816 | 2,944,521,137 | 753 | 0 | 0 |
| Length>=2kb | 2,944,514,837 | 816 | 2,944,521,137 | 753 | 0 | 0 |
| Length>=5kb | 2,944,514,837 | 816 | 2,944,521,137 | 753 | 0 | 0 |

**Supplementary Table S4 Statistics of chromosome-anchored scaffolds of the *C*. *crapnelliana* genome.**

| **Chromosome** | **Length** | **Scaffold Number** |
| --- | --- | --- |
| Chr01 | 235,000,000 | 2 |
| Chr02 | 223,000,000 | 7 |
| Chr03 | 220,000,000 | 12 |
| Chr04 | 212,000,000 | 8 |
| Chr05 | 210,000,000 | 3 |
| Chr06 | 209,000,000 | 3 |
| Chr07 | 198,000,000 | 6 |
| Chr08 | 196,000,000 | 3 |
| Chr09 | 190,000,000 | 5 |
| Chr10 | 184,000,000 | 4 |
| Chr11 | 168,000,000 | 3 |
| Chr12 | 162,000,000 | 6 |
| Chr13 | 154,000,000 | 7 |
| Chr14 | 142,000,000 | 4 |
| Chr15 | 134,000,000 | 5 |
| Total | 2,840,000,000 | 78 |
| **Genome Size** | **2,944,514,837** |  |
| **Chromosome anchoring rate** | **96.34%** |  |

**Supplementary Table S5 BUSCO assessment of the *C. crapnelliana* genome.**

| **Type** | **Number** | **Rate** |
| --- | --- | --- |
| Complete BUSCOs | 1,600 | 99.20% |
| Complete and single-copy BUSCOs | 1,405 | 87.10% |
| Complete and duplicated BUSCOs | 195 | 12.10% |
| Fragmented BUSCOs | 8 | 0.50% |
| Missing BUSCOs | 6 | 0.30% |
| Total Lineage BUSCOs | 1,614 | 100 |

**Supplementary Table S6 BUSCO assessment of gene annotation in the *C*. *crapnelliana* genome.**

| **Type** | **Number** | **Rate (%)** |
| --- | --- | --- |
| Complete BUSCOs | 1,552 | 96.2 |
| Complete and single-copy BUSCOs | 1,409 | 87.3 |
| Complete and duplicated BUSCOs | 143 | 8.9 |
| Fragmented BUSCOs | 22 | 1.4 |
| Missing BUSCOs | 40 | 2.4 |
| Total Lineage BUSCOs | 1,614 | 100 |

**Supplementary Table S7 Statistics of repeat sequences in the *C. crapnelliana* genome.**

| **Class** | **Order** | | **Super family** | **Number of elements** | **Length of sequence (bp)** | **Percentage of sequence (%)** | |
| --- | --- | --- | --- | --- | --- | --- | --- |
| **Class I** |  | |  | 3,578,978 | 1,989,347,501 | 67.56 | |
|  | LINE | |  | 325,084 | 123,372,411 | 4.19 | |
|  |  | | L1 | 49,942 | 20,238,582 | 0.69 | |
|  |  | | Unknown | 265,604 | 99,894,788 | 3.39 | |
|  |  | | RTE-BovB | 9,151 | 3,184,827 | 0.11 | |
|  |  | | Other | 387 | 54,214 | 0 | |
|  | LTR | |  | 3,224,007 | 1862,193,134 | 63.24 | |
|  |  | | Unknown | 1,268,012 | 494,241,207 | 16.79 | |
|  |  | | Copia | 344,938 | 122,547,307 | 4.16 | |
|  |  | | Gypsy | 1,269,154 | 1,070,944,164 | 36.37 | |
|  |  | | Non-chromovirus | 84,021 | 66,237,971 | 2.25 | |
|  |  | | Chromovirus | 29,570 | 12,263,774 | 0.42 | |
|  |  | | Tar | 57,881 | 26,166,593 | 0.89 | |
|  |  | | Ale | 66,839 | 25,153,264 | 0.85 | |
|  |  | | Tork | 38,118 | 14,568,110 | 0.49 | |
|  |  | | Ikeros | 31,327 | 16,714,800 | 0.57 | |
|  |  | | Caulimovirus | 11,963 | 6,128,842 | 0.21 | |
|  |  | | Other | 22,184 | 7,227,102 | 0.25 | |
|  | SINE | |  | 29,887 | 3,781,956 | 0.13 | |
|  |  | | Unknown | 28,352 | 3,616,182 | 0.12 | |
|  |  | | Other | 1,535 | 165,774 | 0.01 | |
| **Class II** |  | |  | 1,018,037 | 327,004,157 | 11.11 | |
|  | DNA | |  | 975,341 | 319,049,746 | 10.84 | |
|  |  | | Unknown | 758,875 | 260,081,367 | 8.83 | |
|  |  | | MULE-MuDR | 40,068 | 10,314,843 | 0.35 | |
|  |  | | hAT-Ac | 40,361 | 10,162,671 | 0.35 | |
|  |  | | CMC-EnSpm | 43,981 | 12,984,980 | 0.44 | |
|  |  | | hAT-Tag1 | 30,967 | 6,296,195 | 0.21 | |
|  |  | | PIF-Harbinger | 13,677 | 3,323,372 | 0.11 | |
|  |  | | MuLE-MuDR | 16,763 | 6,839,086 | 0.23 | |
|  |  | | Other | 30,649 | 9,047,232 | 0.31 | |
|  | MITE | |  | 31,026 | 4,136,861 | 0.14 | |
|  |  | | Unknown | 31,026 | 4,136,861 | 0.14 | |
|  | RC | |  | 11,670 | 3,817,550 | 0.13 | |
|  |  | | Helitron | 11,299 | 3,804,748 | 0.13 | |
|  |  | | Other | 371 | 12,802 | 0 | |
| **Total TEs** |  | |  | **4,597,015** | **2,316,351,658** | **78.67** | |
| **Supplementary Table S7 Statistics of repeat sequences in the *C. crapnelliana* genome (continued).** | | | | | | | |
| **Class** | | **Order** | | **Number of elements** | **Length of sequence (bp)** | | **Percentage of sequence (%)** |
| Tandem Repeats | | |  | 411,936 | 20,392,222 | 0.69 | |
|  | SSR | |  | 220,965 | 2,969,985 | 0.1 | |
|  | Tandem repeat | |  | 190,971 | 17,422,237 | 0.59 | |
| Other |  | |  | 8,580 | 1,751,086 | 0.06 | |
| Unknown |  | |  | 284,599 | 70,683,818 | 2.4 | |
| Simple repeats | | |  | 36,488 | 30,673,669 | 1.04 | |
| Low complexity | | |  | 1,455 | 168,690 | 0.01 | |
| Total Repeats | | |  | 5,340,073 | 2,440,021,143 | 82.87 | |

**Supplementary Table S8 Statistics of non-coding RNA annotation of the *C*. *crapnelliana* genome.**

| **Type** |  | **Number** | **Average_length (bp)** | **Total_length (bp)** | | **Percentage (%)** |
| --- | --- | --- | --- | --- | --- | --- |
| **Regulatory** |  | 45 | 48.2 | 2,169 | 0.0001 | |
| **tRNA** |  | 857 | 76.5 | 65,560 | 0.0022 | |
| **rRNA** | **rRNA** | 7,988 | 1,438.31 | 11,489,216 | 0.3902 | |
|  | **18S** | 1,508 | 1,813.63 | 2,734,955 | 0.0929 | |
|  | **28S** | 1,477 | 5,504.41 | 8,130,012 | 0.2761 | |
|  | **5S** | 4,974 | 117.32 | 583,543 | 0.0198 | |
|  | **5.8S** | 16 | 153.69 | 2,459 | 0.0001 | |
| **ncRNA** | **ncRNA** | 838 | 124.01 | 103,923 | 0.0035 | |
|  | **other** | 46 | 282.7 | 13,004 | 0.0004 | |
|  | **snRNA** | 485 | 103.84 | 50,360 | 0.0017 | |
|  | **miRNA** | 176 | 128.07 | 22,541 | 0.0008 | |
|  | **spliceosomal** | 131 | 137.54 | 18,018 | 0.0006 | |

**Supplementary Table S9 Statistics of functional annotation of protein-coding genes in the *C. crapnelliana* genome.**

| **Item** | **Count** | **Percentage** |
| --- | --- | --- |
| All | 37,390 | 100.00% |
| Annotation | 37,015 | 99.00% |
| KOG | 20,611 | 55.12% |
| KEGG | 14,438 | 38.61% |
| NR | 36,997 | 98.95% |
| SwissProt | 30,236 | 80.87% |
| GO | 23,555 | 63.00% |

**Supplementary Table 10 Summary of gene annotation in the *C. crapnelliana* and other representative plant genomes.**

| **Species** | **Number of gene** | **Average gene length**  **(bp)** | **Average CDS length**  **(bp)** | **Average exons per gene** | **Average exon length**  **(bp)** | **Average introns per gene** | **Average intron length**  **(bp)** |
| --- | --- | --- | --- | --- | --- | --- | --- |
| ***C. crapnelliana*** | 37,390 | 9,590.45 | 1,254.23 | 5.54 | 226.29 | 4.54 | 1,835.12 |
| *Actinidia chinensis* | 42,953 | 4,800.67 | 1,005.82 | 5.06 | 198.96 | 4.06 | 935.78 |
| *Arabidopsis thaliana* | 27,412 | 1,856.57 | 1,205.21 | 5.09 | 236.78 | 4.09 | 159.26 |
| *Beta vulgaris* | 35,855 | 3,526.68 | 933.61 | 3.66 | 255.36 | 2.66 | 976.27 |
| *Brachypodium distachyon* | 34,309 | 2,501.50 | 1,119.58 | 4.41 | 253.81 | 3.41 | 405.11 |
| *Camellia impressinervis* | 42,499 | 4,904.76 | 1,187.71 | 5.02 | 236.58 | 4.02 | 924.56 |
| *Camellia sinensis* | 49,411 | 5,511.19 | 1,080.38 | 4.89 | 220.97 | 3.89 | 1,139.25 |
| *Coffea canephora* | 25,574 | 3,188.40 | 1,205.55 | 5.1 | 236.22 | 4.1 | 483.2 |
| *Diospyros lotus* | 23,977 | 8,500.09 | 1,341.31 | 5.44 | 246.56 | 4.44 | 1,612.26 |
| *Manihot esculenta* | 33,034 | 2,962.18 | 1,160.45 | 4.83 | 240.12 | 3.83 | 470.08 |
| *Olea europaea* | 39,716 | 3,685.61 | 1,163.21 | 4.77 | 243.93 | 3.77 | 669.31 |
| *Oryza sativa* | 55,974 | 2,569.13 | 1,372.74 | 4.1 | 334.65 | 3.1 | 385.68 |
| *Rhododendron delavayi* | 32,938 | 3,993.95 | 1,153.21 | 4.62 | 249.7 | 3.62 | 785.08 |
| *Rosa chinensis* | 30,767 | 2,951.57 | 1,330.84 | 4.94 | 269.37 | 3.94 | 411.3 |
| *Solanum lycopersicum* | 25,476 | 4,093.47 | 1,303.10 | 5.35 | 243.49 | 4.35 | 641.2 |
| *Vitis vinifera* | 25,538 | 5,709.29 | 1,337.71 | 5.2 | 257.12 | 4.2 | 1,040.21 |

**Supplementary Table S11 Orthologous groups identified in the *C. crapnelliana* and other plant genomes.**

| **Species** | **Unclustered orthologs** | **Unique paralogs** | **Single-copy orthologs** | **Multiple-copy orthologs** |
| --- | --- | --- | --- | --- |
| *Vitis vinifera* | 7,213 | 6,750 | 6,472 | 41,879 |
| *Arabidopsis thaliana* | 3,130 | 6,622 | 6,276 | 25,980 |
| *Olea europaea* | 5,012 | 11,729 | 6,624 | 39,048 |
| *Actinidia chinensis* | 1,144 | 590 | 5,763 | 26,208 |
| *C. oleifera* | 1,700 | 4,121 | 6,628 | 38,284 |
| *C. chekiangoleosa* | 3,593 | 4,438 | 10,101 | 50,690 |
| ***C. crapnelliana*** | 1,603 | 837 | 10,169 | 25,618 |
| *C. sinensis* var. *assamica* | 3,129 | 1,248 | 10,396 | 35,371 |
| *C. sinensis* var. *sinensis* | 2,735 | 794 | 10,506 | 29,373 |

**Supplementary Table S12 The key gene families involved in the fatty acid biosynthesis in *A. thaliana*.**

| **Pathway** | **Protein Family** | **Protein Family Abbreviation** | **pfam.**  **name** | **pfam.id** |
| --- | --- | --- | --- | --- |
| Fatty Acid Synthesis | Carboxyltransferase alpha Subunit of Heteromeric ACCase | alpha-CT | ACCA | PF03255.14 |
| Fatty Acid Synthesis | Carboxyltransferase beta Subunit of Heteromeric ACCase | beta-CT | Carboxyl_trans | PF01039.22 |
| Fatty Acid Synthesis | Biotin Carboxylase; subunit of Heteromeric ACCase | BC | Biotin_carb_C | PF02785.19 |
|  |  | BC | Biotin_carb_N | PF00289.22 |
|  |  | BC | CPSase_L_D2 | PF02786.17 |
| Fatty Acid Synthesis | Biotin Carboxyl Carrier Protein; subunit of Heteromeric ACCase | BCCP | Biotin_lipoyl | PF00364.22 |
| Fatty Acid Synthesis | Ketoacyl-ACP Synthase I | KASI | ketoacyl-synt | PF00109.26 |
|  |  | KASI | Ketoacyl-synt_C | PF02801.22 |
| Fatty Acid Synthesis | Ketoacyl-ACP Synthase II | KASII | ketoacyl-synt | PF00109.26 |
|  |  | KASII | Ketoacyl-synt_C | PF02801.22 |
| Fatty Acid Synthesis | Ketoacyl-ACP Synthase III | KASIII | ACP_syn_III_C | PF08541.10 |
|  |  | KASIII | ACP_syn_III | PF08545.10 |
| Fatty Acid Synthesis | Acyl-ACP Thioesterase A | FATA | Acyl-ACP_TE | PF01643.17 |
| Fatty Acid Synthesis | Acyl-ACP Thioesterase B | FATB | Acyl-ACP_TE | PF01643.17 |
|  |  | FATB | Acyl-thio_N | PF12590.8 |
| Fatty Acid Synthesis | Stearoyl-ACP Desaturase | SAD | FA_desaturase_2 | PF03405.14 |
| Triacylglycerol Biosynthesis | Oleate Desaturase | FAD2-8 | FA_desaturase | PF00487.24 |
| Prokaryotic Galactolipid | Linoleate Desaturase | FAD4 | TMEM189_B_dmain | PF10520.9 |
| Triacylglycerol Biosynthesis | Oil-Body Oleosin | OBO | Oleosin | PF01277.17 |
| Triacylglycerol Biosynthesis | Caleosin | CALO | Caleosin | PF05042.13 |
| Triacylglycerol Biosynthesis | Steroleosin | STERO | Steroleosin | PF00106.26/PF13561.7 |

**Supplementary Table S13 Copy number of candidate genes involved in the fatty acid biosynthesis in *C. crapnelliana*, tea tree and *A. thaliana* genomes.**

| **Gene family** | ***Arabidopsis thaliana*** | ***C. sinensis* var. *assamica*** | ***C. crapnelliana*** |
| --- | --- | --- | --- |
| ***α-CT*** | 1 | 4 | 4 |
| ***β-CT*** | 1 | 5 | 3 |
| ***BC*** | 1 | 2 | 4 |
| ***BCCP*** | 2 | 3 | 3 |
| ***KASI/KASII*** | 2 | 9 | 6 |
| ***KASIII*** | 1 | 2 | 4 |
| ***FATA*** | 2 | 4 | 5 |
| ***FATB*** | 1 | 3 | 3 |
| ***SAD*** | 7 | 12 | 8 |
| ***FAD2*** | 1 | 2 | 2 |
| ***FAD3*** | 1 | 1 | 2 |
| ***FAD4*** | 3 | 1 | 1 |
| ***FAD5*** | 9 | 1 | 3 |
| ***FAD6*** | 1 | 1 | 7 |
| ***FAD7*** | 1 | 0 | 0 |
| ***FAD8*** | 1 | 4 | 2 |
| **Total** | **35** | **54** | **57** |
| ***OBO*** | 8 | 9 | 5 |
| ***CALO*** | 8 | 4 | 5 |
| ***STERO*** | 8 | 124 | 104 |
| **Total** | **24** | **137** | **114** |
